# Strengthening intraguild predation increases temporal variability of biomass across trophic levels in model food webs

**DOI:** 10.1101/2025.06.25.661600

**Authors:** Chase J. Rakowski, Mathew A. Leibold, Caroline E. Farrior

**Affiliations:** Department of Integrative Biology, University of Texas at Austin, Austin, TX, USA; Institute of Arctic and Alpine Research, University of Colorado at Boulder, Boulder, CO, USA; Department of Biology, University of Florida, Gainesville, FL, USA; Institut Natura e Teoria en Pirineus, 09400, Surba, France

**Keywords:** ecosystem functioning, ecosystem stability, IGP, interaction strength, multitrophic, omnivory, plankton, population dynamics, temporal variability, top-down control, trophic cascade

## Abstract

Multiple global-change forces, from habitat alterations to warming, are altering food webs and trophic interaction strengths. Such changes in trophic interactions have important implications, as it is a tenet of ecology that trophic interactions are linked to the functioning and stability of ecosystems. For example, changes in the presence or strength of intraguild predation (IGP), the consumption of a predator by another predator that competes for shared prey, can have cascading effects on the biomasses of species and trophic levels. For this reason, IGP can affect key ecosystem functions at the base of the food web and is of special interest to practitioners of biological pest control. However, the relationship between IGP and ecosystem stability is not yet well understood, especially whether and how IGP might affect the stability of non-adjacent lower trophic levels including primary producers. In this study we simulate the dynamics of a six-species, four-trophic-level food web plus a limiting nutrient to explore the relationship between IGP strength and the temporal variability of species- and trophic group-biomass. By varying the IGP rate given the abundance of the eaten predator, we find that the model food web abruptly shifts between equilibria in which all species maintain either constant biomass or stable limit cycles where all trophic levels exhibit sustained and significant oscillations. While complex feedback in the model creates a divergence between the IGP functional response and the resulting realized IGP strength, both stronger IGP functional responses and stronger realized IGP are associated with a higher likelihood of oscillations. Furthermore, analyses indicate that the strongest consumptive interaction induces the oscillating behavior in an indirect effect initiated by the change in IGP. Overall, these results suggest that as food web structure changes in ecosystems worldwide, strengthening IGP runs the risk of inducing destabilizing effects that extend to the base of food webs, while weakening IGP could confer stability to ecosystem functions such as primary production. Finally, we discuss relevance to management, including the implication that IGP among biological control agents should be minimized to maintain stable crop production.

## INTRODUCTION

Global change forces are altering community composition and predation rates in ecosystems around the world, thereby changing food web structure in local communities worldwide. Predators face fast extinction rates yet are sometimes replaced by introduced predator species, while other aspects of global change are modifying relative species abundances, leading to changes in predator and prey composition (Purvis et al. 2000, Cucherousset et al. 2012, Bond and Lavers 2014, Hu et al. 2022). Furthermore, aspects of global change such as warming can directly and indirectly affect predation rates independent of changes in community composition (Hammill et al. 2018, Frances and McCauley 2018). For these reasons trophic interaction strengths are likely to change in many ecosystems (Frances and McCauley 2018). One such trophic interaction prone to change is intraguild predation (IGP), a very common aspect of food webs defined as the consumption of a predator by another predator which competes for shared prey (Polis et al. 1989).

Food web structure is generally an important determinant of ecosystem functioning, and IGP appears to be no exception. Species diversity, the prevalence of weak trophic interactions and omnivory, and the number of trophic levels can all affect ecosystem functions and stability (McCann and Hastings 1997, McCann et al. 1998, Thébault and Loreau 2005, 2006, Jiang and Pu 2009, Tilman et al. 2014). IGP in particular has been shown to affect the biomass of various trophic levels. Specifically, empirical and theoretical work has shown that IGP can often weaken total top-down control by multiple predator species, and this effect can reverberate through trophic levels to alter trophic cascades (Finke and Denno 2004, Straub et al. 2008). Additionally, IGP can have varying effects on trophic level biomasses depending on its strength (Chang et al. 2020, Chang and Cardinale 2020).

The nature of the relationship between IGP and ecosystem stability is murkier. Relevant empirical investigations are rare, and theoretical studies offer a mixture of conclusions. Some papers suggest that IGP or omnivory – a broader category including IGP – is generally destabilizing, both in the sense of constraining coexistence and in the sense of creating oscillations and chaos (Pimm and Lawton 1978, Holt and Polis 1997, Tanabe and Namba 2005, Hossain et al. 2020). However, other studies emphasize that omnivory can either destabilize or stabilize depending on omnivory strength and other conditions (McCann and Hastings 1997, Vandermeer 2006), and Křivan (2000) found that IGP strength is positively correlated with stability. Furthermore, these studies analyzed models with only three species, limiting the complex food web feedbacks and prey switching that could underlie any relationship between trophic interaction strength and stability in more diverse food webs (Post et al. 2000). Indeed, adding alternative prey species to the three-species module enables a richer menu of dynamical outcomes (P. Daugherty et al. 2007, Holt and Huxel 2007). For example, in versions of these models with saturating functional responses, adding weak predator-prey interactions stabilizes the system while adding strong interactions destabilizes it (Holt and Huxel 2007). However, these analyses of more diverse IGP food web models did not explicitly explore the relationship between IGP and stability, and in particular, it is not yet known whether IGP strength might influence the variability of biomass in non-adjacent lower trophic levels. Such a relationship would be especially important because IGP commonly occurs among predators multiple trophic levels above primary producers; yet, primary production provides the foundation of ecosystems, climate regulation, and crop production. Therefore, a link between IGP and the stability of basal trophic levels would carry important implications both for natural ecosystems and for biological control of agroecosystems involving multiple natural enemy species, a potential link involving four trophic levels (top predator ➔ mesopredator ➔ herbivore ➔ plant) which smaller modules cannot replicate. Extending IGP models to more complete food webs is not a trivial extension because additional species create the possibility of complex feedback that can lead to unexpected emergent results.

Here we use a six-species, four-trophic level food web model (two primary producers, two herbivores, and a smaller predator consumed by a top predator) to test the effects of IGP on the central tendency of biomass and its variability throughout the food web. To explore a range of IGP strengths, we manipulate the maximum IGP rate, a parameter that defines the rate of IGP given density of the smaller predator, rather than adding or removing IGP altogether. The maximum IGP rate captures an aspect of the top predator’s behavior akin to its handling time or preference for the smaller predator relative to other prey species. However, this parameter is not the same as the realized IGP strength, defined here as biomass of the smaller predator consumed per unit time per unit biomass of the top predator, which depends on the maximum IGP rate as well as the densities of all the predator and prey species. We predicted that weaker IGP (of either definition) would allow greater total top-down control of the herbivorous shared prey, leading to reduced herbivore biomass and that the effect would extend to a trophic cascade, with reduced herbivory allowing producers to maintain higher and more stable biomass. Interestingly, we found that the maximum IGP rate has a complex relationship with realized IGP strength due to the feedback among species. Changes in the maximum IGP rate cause state changes and complex rearrangements in biomasses across the food web, thereby altering interaction strengths including IGP strength in a nonlinear manner. However, in general both lower maximum IGP rates and weaker realized IGP are associated with stable biomass of all species and trophic levels.

## METHODS

We simulate temporal dynamics of food webs with a single limiting nutrient *N*, two primary producer species *A*_1_ and *A*_2_, two herbivorous species *H*_1_ and *H*_2_ which consume both *A*_1_ and *A*_2_, a predator species *P*_1_ that consumes both *H*_1_ and *H*_2_, and a top predator *P*_2_ that consumes both *H*_1_ and *H*_2_ as well as *P*_1_, representing IGP (Fig. 1a). Within a trophic group, the species with subscript 1 is smaller-bodied than the species with subscript 2, which affects their growth rates and other relevant properties via allometric scaling as explained below. Our model is based on the work by Ceulemans et al. (2019) who study relatively complex food web dynamics in a plankton chemostat framework but exclude IGP. Besides adding IGP and making some relevant parameter changes (see below), we also reduce the model to two rather than four herbivore species, and alter the model to represent natural food webs by allowing mortality rates to vary across species.

**Figure 1:**
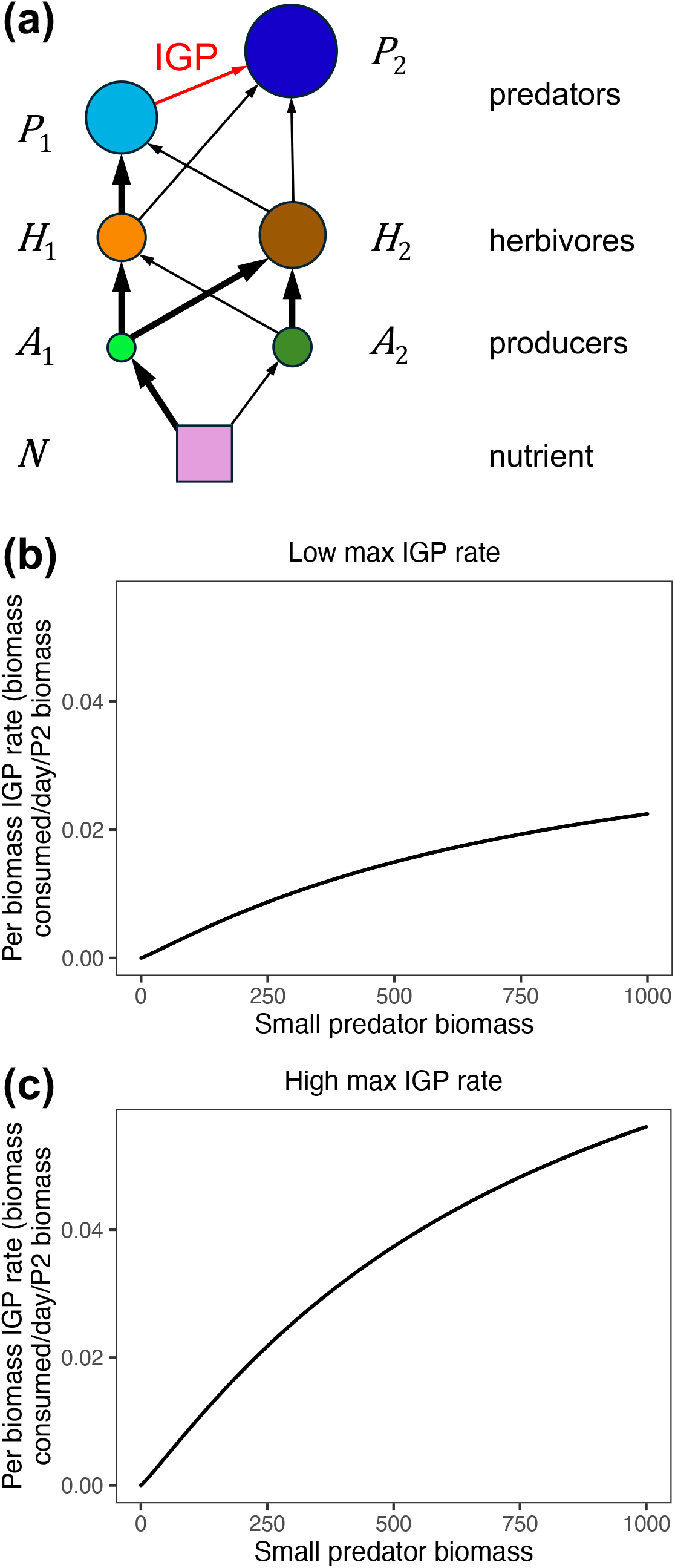
**a)** Diagram representing the model food web structure. Circles represent species; relative circle sizes represent qualitative differences in body size. Arrows represent energy flow, with thickness representing qualitative differences in relative per capita consumption rates (determined by half-saturation constants 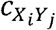 and maximum consumption rates *r*_*Y*_) within trophic levels. The red arrow represents intraguild predation (IGP). **b-c:** Example functional response curves showing IGP rate over density (biomass) of the small predator *P*_1_, which in this context is the prey. **b)** 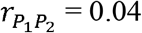 (low IGP rate at high *P*_1_ densities); **c)** 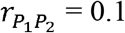 (high IGP rate at high *P*_1_ densities).

We simulate biomass changes in each species from one time point to the next as a function of its own abundance, the abundance of its predators, and the abundance of its resources through time. The concentration of available *N, n*, is replenished at a rate *δ* proportional to its current deviation from the loading *N* concentration *n*_0_. *N* is taken up by the producers (*A*_1_, *A*_2_) following a Monod function:

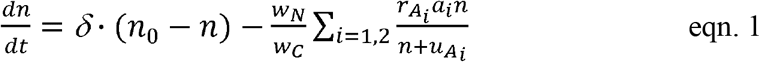

where 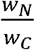 is the *N*:carbon weight ratio of producers (parameterized for nitrogen), 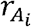 is the intrinsic rate of increase of *A*_*i*_, *a*_*i*_ is the biomass of *A*_*i*_, and *u*_*i*_ is the half-saturation constant of *N* uptake by *A*_*i*_.

We model the consumption of a prey species at any trophic level *X* by a predator species, *Y*, with the functional response:

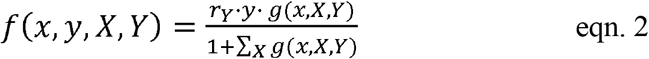

where *r*_*Y*_ is the maximum consumption rate of the predator *Y*, equivalent to the inverse of handling time, *y* is the biomass of *Y, x* is the biomass of *X*, and

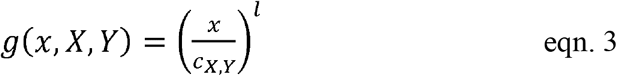

where *l* controls the curvature of the functional response at low prey densities (*l* =1 yields a Holling type II while *l* =2 yields a Holling type III), and *c*_*X,Y*_ is the half-saturation constant of the consumption of *X* by *Y*. These equations apply to any consumption of another species including IGP, in which case *r*_*Y*_ is 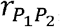, *X* is *P*_1_ in the numerator and *H*_1_, *H*_2_, and *P*_1_ in the denominator, and *Y* is *P*_2_ (see eqn. 8 below). The Hill exponent *l* is primarily set to 1.1 so that the functional response is saturating but is less destabilizing than Holling type II (Fig. 1b-c; Williams and Martinez 2004), although we also conducted a sensitivity analysis with respect to *l* (methods below). When *c*_*X,Y*_ is high, *Y* consumes *X* at a low rate at a given biomass density of *X* (*x*), and vice versa. Conversely, when *r*_*Y*_ is high, *Y* consumes *X* at a high rate at a given biomass density of *X* (*x*), and vice versa. Note that consumption of one prey species, *X*_1_ is influenced by the other prey species, *X*_2_ indirectly here due to predator satiation captured by the sum over all prey species in eqn. 2.

The change in biomass of producer species *A*_*i*_ (*a*_*i*_) is governed by:

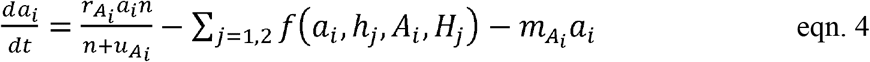

where *h*_*j*_ represents the biomass of *H*_*j*_ and *m*_*Ai*_ is the baseline mortality rate of *A*_*i*_ apart from herbivory. The change in biomass of a heterotroph species (*Y*_*j*_) that consumes *X* species and is consumed by *Z* species is:

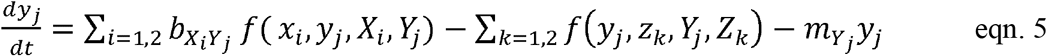

where 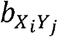 is the conversion efficiency of *X*_*i*_ to *Y*_*j*_ and 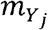 is the baseline mortality rate of *Y*_*j*_. For herbivores (*H* species), *A* species take the place of the *X*’s and *P’s* take the place of the *Z’s*. For *P*_1_, *X* = {*H*_1_, *H*_2_} and *Z* = *P*_2_. The change in *p*_2_ is:

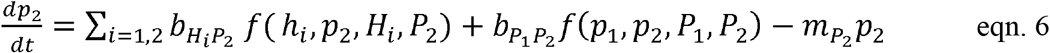

where *X* = {*H*_1_, *H*_2_, *P*_1_}. The functional responses of predators for given prey items, *f*, are given by eqn. 2. Parameters *r* and *m* are different for each *Y*_*j*_, and *c* is different for each combination of *X*_*i*_ and *Y*_*j*_. The conversion efficiency *b* is kept the same except for the case of IGP, where it is increased (see below). All differential equations that govern the model are written in Appendix S1 (eqns. S1-S5).

We primarily use parameters from Ceulemans et al. (2019) but with several exceptions. While Ceulemans et al. (2019) varied within-trophic level species trait diversity, we only use the parameters representing 75% of the maximal species trait diversity used in their study for pairs of species in a trophic group. The parameters vary based on allometric scaling (Table 1):

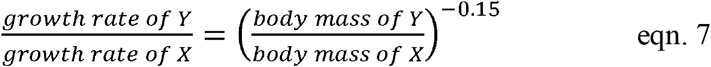

where, as in Ceulemans et al. (2019), we assume body mass ratios of 10^3^ across adjacent trophic levels and an allometric scaling exponent of -0.15. However, rather than setting all mortality rates equal to each other to simulate a chemostat experiment, we use the same allometric scaling exponent to determine mortality rates across species and better represent natural food webs.

**Table 1:** Central parameter values used in the model.

| Parameter | Value | Description |
| --- | --- | --- |
| $\delta$ | 0.055 | available nutrient loss and replenishment rate |
| $n_0$ | 1120 $\mu\text{gN/L}$ | incoming/initial concentration of available nutrient |
| $\frac{w_N}{w_C}$ | 0.175 $\text{gN/gC}^*$ | nutrient:carbon ratio of $A_i$ |
| $u_{A_1}$ | 8 $\mu\text{gN/L}$ | half-saturation constant for nutrient uptake by $A_1$ |
| $u_{A_2}$ | 13 $\mu\text{gN/L}$ | half-saturation constant for nutrient uptake by $A_2$ |
| $r_{A_1}$ | 0.95/day | intrinsic rate of increase of $A_1$ |
| $r_{A_2}$ | 0.68/day | intrinsic rate of increase of $A_2$ |
| $r_{H_1}$ | 1.01/day | maximum herbivory rate by $H_1$ (inverse of handling time) |
| $r_{H_2}$ | 0.73/day | maximum herbivory rate by $H_2$ (inverse of handling time) |
| $r_{P_1}$ | 0.38/day | maximum predation rate by $P_1$ (inverse of handling time) |
| $r_{P_2}$ | 0.06/day | maximum herbivore predation rate by $P_2$ (inverse of handling time) |
| $r_{P_1P_2}$ | varied | maximum IGP rate (consumption of $P_1$ by $P_2$; inverse of handling time) |
| $c_{A_1H_1}$ | 300 $\mu\text{gC/L}$ | half-saturation constant for consumption of $A_1$ by $H_1$ |
| $c_{A_2H_1}$ | 850 $\mu\text{gC/L}$ | half-saturation constant for consumption of $A_2$ by $H_1$ |
| $c_{A_1H_2}$ | 400 $\mu\text{gC/L}$ | half-saturation constant for consumption of $A_1$ by $H_2$ |
| $c_{A_2H_2}$ | 400 $\mu\text{gC/L}$ | half-saturation constant for consumption of $A_2$ by $H_2$ |
| $c_{H_1P_1}$ | 400 $\mu\text{gC/L}$ | half-saturation constant for consumption of $H_1$ by $P_1$ |
| $c_{H_2P_1}$ | 650 $\mu\text{gC/L}$ | half-saturation constant for consumption of $H_2$ by $P_1$ |
| $c_{H_1P_2}$ | 650 $\mu\text{gC/L}$ | half-saturation constant for consumption of $H_1$ by $P_2$ |
| $c_{H_2P_2}$ | 550 $\mu\text{gC/L}$ | half-saturation constant for consumption of $H_2$ by $P_2$ |
| $c_{P_1P_2}$ | 800 $\mu\text{gC/L}$ | half-saturation constant for IGP (consumption of $P_1$ by $P_2$) |
| $l$ | 1.1 | Hill exponent (controls consumption rate at low prey densities) |
| $b_{A_iH_j}$ | 0.33* | conversion efficiency of $A_i$ into new $H_j$ biomass |
| $b_{H_jP_k}$ | 0.33* | conversion efficiency of $H_j$ into new $P_k$ biomass |
| $b_{P_1P_2}$ | 0.6 | conversion efficiency of $P_1$ into new $P_2$ biomass |
| $m_{A_1}$ | 0.075 | density-independent mortality rate of $A_1$ |
| $m_{A_2}$ | 0.0485 | density-independent mortality rate of $A_2$ |
| $m_{H_1}$ | 0.04 | density-independent mortality rate of $H_1$ |
| $m_{H_2}$ | 0.03 | density-independent mortality rate of $H_2$ |
| $m_{P_1}$ | 0.01 | density-independent mortality rate of $P_1$ |
| $m_{P_2}$ | 0.01 | density-independent mortality rate of $P_2$ |

\*Not varied in the sensitivity analyses (see Methods).

Species in lower trophic levels have lower body masses, and one species is smaller than the other within each trophic level. Smaller species have higher maximum growth rates 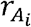 for producers and 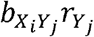 for consumers), higher rates of nutrient uptake or prey consumption (lower 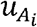 or lower 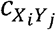: see eqn. 1 and 3, respectively), and higher baseline mortality rates 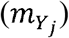, but they are less defended – that is, they are consumed faster. Within trophic levels, we assume the smaller consumer species consumes the smaller prey species at a higher rate (lower 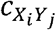) than it consumes the larger prey species, whereas the larger herbivore is a generalist, which is to say 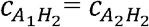. While Ceulemans et al. (2019) modeled four herbivorous species representing all possible combinations of the defended/undefended and selective/generalist traits, we model only an undefended selective (*H*_1_) and a defended generalist herbivore (*H*_2_) to simplify the analysis and focus on the case where the herbivores have a tradeoff with the best chance of enabling coexistence.

To allow exploration of a range of IGP strength, we alter some of the predators’ parameters from their allometrically-determined values in a way that enables the predators to coexist in the presence of IGP. First, we assume the predators have a tradeoff such that IGP is especially beneficial for *P*_2_ (higher conversion efficiency, 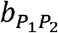, than for other interactions), but that the smaller predator *P*_1_ is a superior competitor for their shared herbivore prey (lower 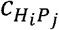 value; Holt and Polis 1997, P. Daugherty et al. 2007). Additionally, we lowered *P*_2_’s maximum herbivore predation rate 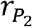. Finally, the predators’ baseline mortality rates are set equal to each other rather than setting *P*_1_’s mortality rate higher in line with its smaller size (Table 1). Note that the smaller predator’s effective mortality rate, including both baseline mortality and mortality due to predation, is still higher than that of the top predator. Also note that, as with other parameters, the predators’ mortality rates were separately varied and therefore no longer equal in the sensitivity analysis described below.

We ran simulations across the entire range of values of the maximum IGP rate 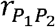 for which all 6 species coexist. The parameter 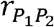 is equivalent to the inverse of handling time for the IGP interaction and along with other parameters, especially 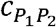, it determines the degree to which *P*_2_ exploits *P*_1_ at a given density of *P*_1_. Species extinction was defined by the biomass of a species falling below 0.01 µgC/L after removing transient dynamics. To test the robustness of our results, we re-ran this parameter sweep of 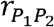 for different sets of parameter values for nearly all other parameters (Table 1). Each set of parameters was developed by randomly and separately varying parameters by up to 10% above and 10% below the central value, then retaining parameter combinations that adhered to the allometric assumptions. These assumptions include that compared to larger species, smaller species grow faster, take up nutrients or consume prey faster, and experience faster baseline mortality rates (except the smaller vs. top predator), and that within trophic groups, the smaller species consume smaller prey faster than they consume larger prey (see Appendix S1: Table S1 for specific parameter constraints). We also conducted separate sensitivity analyses for nutrient enrichment, *n*_0_, and the Hill exponent, *l*, by varying these parameters by up to 10% above and 10% below their central values while varying initial predator biomasses as in the main simulation analyses but keeping all other parameters constant.

In all cases, we ran simulations for each parameter combination until equilibrium, either a fixed point or a stable limit cycle, was reached. Then we removed transient dynamics and recorded available nutrient masses, species biomasses, and trophic interaction strengths (nutrient uptake rates per unit biomass of the producer, and consumption rates per unit biomass of the consumer) for at least 2,500 days (6.85 years). Importantly, the realized trophic interaction strengths are different from the parameters defining trophic interactions. For example, while we manipulated the maximum IGP rate 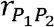, we measured the realized IGP strength as the biomass of *P*_1_ consumed per day per unit biomass of *P*_2_ as defined in the model (eqns. 2 and 5, Appendix S1 eqn. S4):

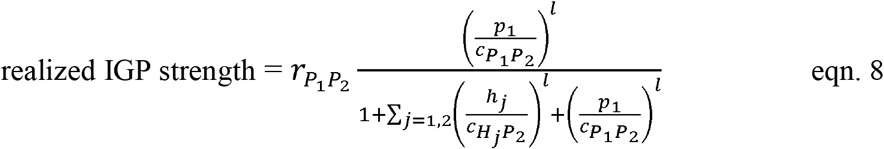

where *p*_1_, *h*_1_, and *h*_2_, are results of the simulation.

To test the effects of initial conditions on long-run biomass dynamics, we initiated simulations with various species biomasses. We first initiated simulations with all species at 500 µg/L biomass. We then re-initiated simulations with each species, one at a time, beginning at 1% of this value (5 µg/L) and again beginning at 2500—which is near the highest constant species biomass observed in simulations—while the other species were initiated at 500. Then we repeated the trials for pairs of species, simultaneously varying the initial biomass of both producers, both herbivores, and both predators, using combinations of the same initial biomasses of 5, 500, and 2500. We also repeated these trials beginning all other species at their median long-run biomasses instead of 500. All aforementioned trials were completed for each value of 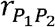 within the central parameter set and the first three randomized parameter sets. Upon finding that switching among alternative stable states was associated with the predators’ initial biomasses, we also conducted more fine-grained trials initiating the predators at combinations of 50 biomasses between 0.5 and 10,000 at 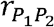 values near state transitions to ensure we observed the full extent of alternative stable states across 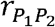 values. In all cases, all possible alternative stable states were reached across initial predator biomasses of {*P*_1_ = 5, *P*_2_ = 2500}, {*P*_1_ = 500, *P*_2_ = 500}, and {*P*_1_ = 2500, *P*_2_ = 5}, with other species initiated at 500. Therefore, we used these starting conditions to analyze all subsequent randomized parameter sets.

We further conducted analyses to better understand which species induce biomass oscillations. To do so, we first ran simulations with 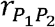 values that cause an oscillating state, except that we forced the biomass of one individual species to remain constant at its median. Then we repeated this exercise with every other species, and in both oscillating states (1 and 3), allowing us to identify which species are responsible for the oscillations. These analyses were conducted using the central parameter values (Table 1).

All simulations and analyses were conducted in R v. 4.4.0 (R Core Team 2022).

## RESULTS

Our results show that there is a complex, up to three-phase effect of varying the maximum IGP rate 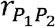 —which determines the rate of IGP at given densities of the predators— in which the food web abruptly transitions between sustained oscillations and point equilibria (Fig. 2). Note that under randomized parameter sets, simpler two-phase relationships with a point equilibrium and an oscillating equilibrium were also common (Appendix S1: Fig. S1). The point equilibria are generally associated with low 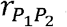 and low realized IGP strength (eqn. 8).

**Figure 2:**
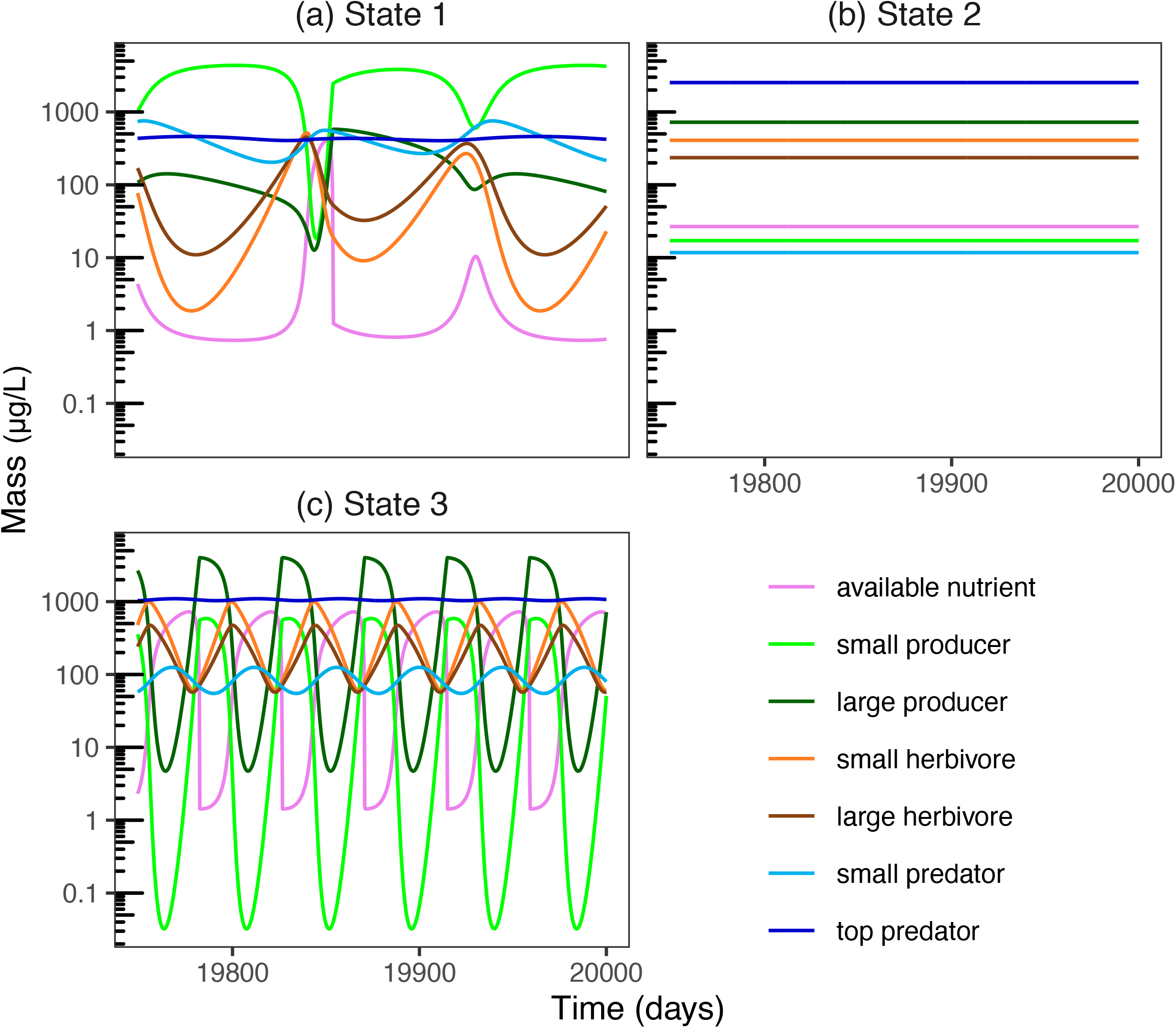
Example time series demonstrating long-run behavior of masses of each species and of the available nutrient for each of the three qualitative states. **a)** State 1, slow oscillations 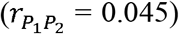: **b)** State 2, static 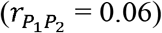: **c)** State 3, fast oscillations 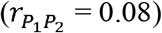. All simulations were run with the central parameters (Table 1).

However, we also find qualitatively distinct oscillations at very low versus high 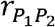, both of which are associated with high realized IGP strength. Therefore, the maximum IGP rate 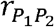 and the realized IGP strength are often unaligned. Furthermore, the transitions from stability to both oscillation types often depend on initial predator biomasses. Adding to the complexity of results, we find that for some values of low and high 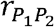, one or more species go extinct under some initial predator biomasses while a stable coexisting state is reached under different initial predator biomasses (Fig. 3, Appendix S1: Fig. S1).

**Figure 3:**
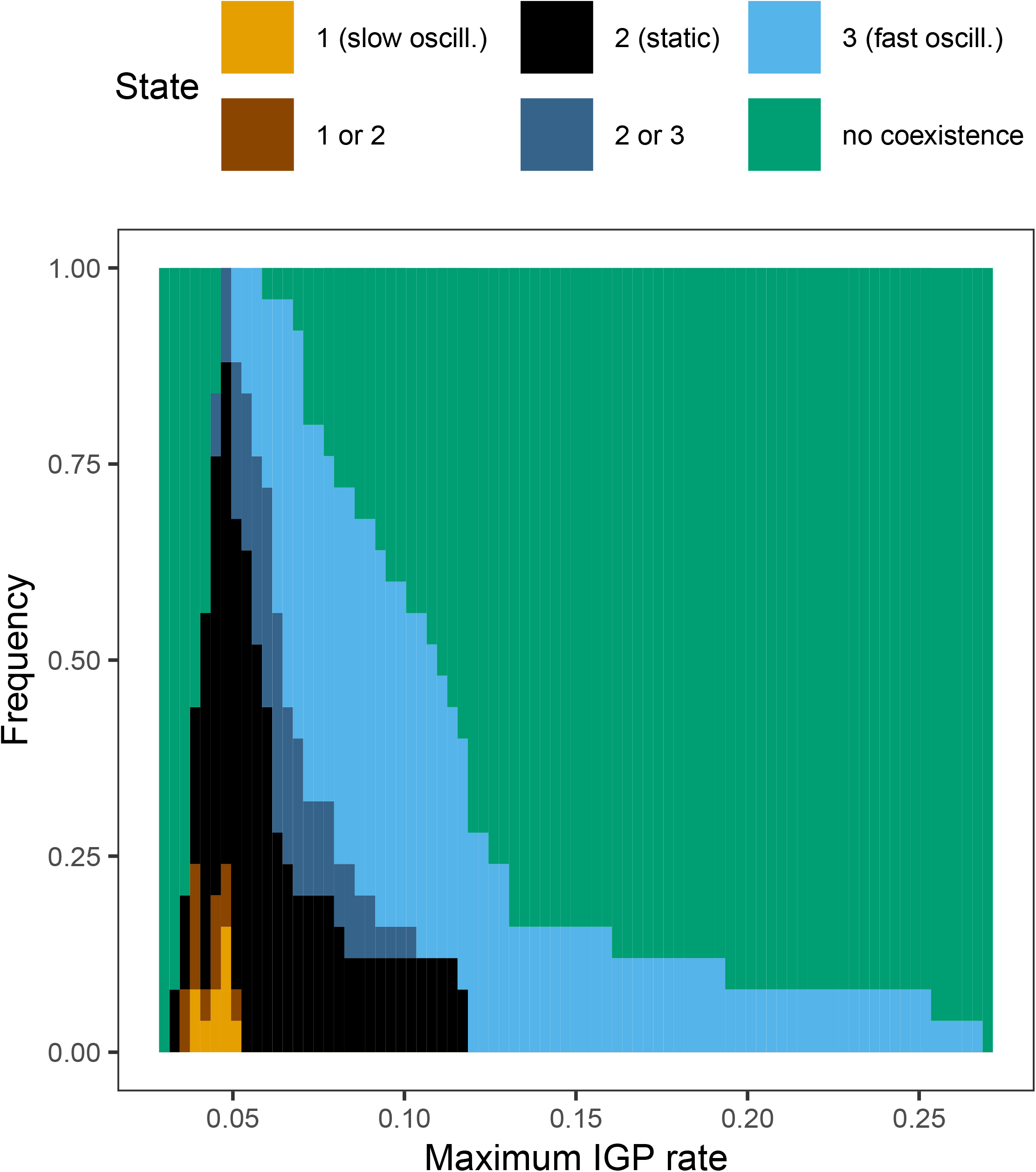
Stacked bar chart representing the frequency of qualitative equilibrium states across maximum IGP rates 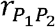, summarized across all randomized parameter sets. Possible states include State 1: slow oscillations, State 2: point equilibrium, and State 3: fast oscillations, as well as alternative stable states where State 1 or 2 are possible or where State 2 or 3 are possible, depending on initial conditions, and finally the outcome where all six species fail to coexist (“no coexistence”).

### Qualitative states

State 1 is characterized by a stable limit cycle of slow biomass oscillations and high producer biomass, specifically a very high *A*_1_ biomass despite *A*_2_ biomass being very low. The available nutrient is quickly taken up by the large *A*_1_ population and does not often reach a high concentration (Fig. 4a, Appendix S1: Figs. S2a, S3). Herbivore and predator biomass is also somewhat low, although *P*_1_ stays at a higher biomass than in the other states, uniquely resulting in similar biomass of the two predators. State 1 is the equilibrium for a small range of very low maximum IGP rates: 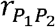 between 0.043 and 0.045 with the central parameters (Fig. 4, Appendix S1: Fig. S3), and between 0.036 and 0.051 among all randomized parameter combinations studied (Fig. 3, Appendix S1: Fig. S1). Under some parameter combinations State 1 is never possible (Fig. 3, Appendix S1: Fig. S1). Under the central parameters, the system only reaches State 1 if both predators begin with moderate biomasses, e.g. 500 each; if one predator begins with a much higher biomass than the other, it reaches State 2.

**Figure 4:**
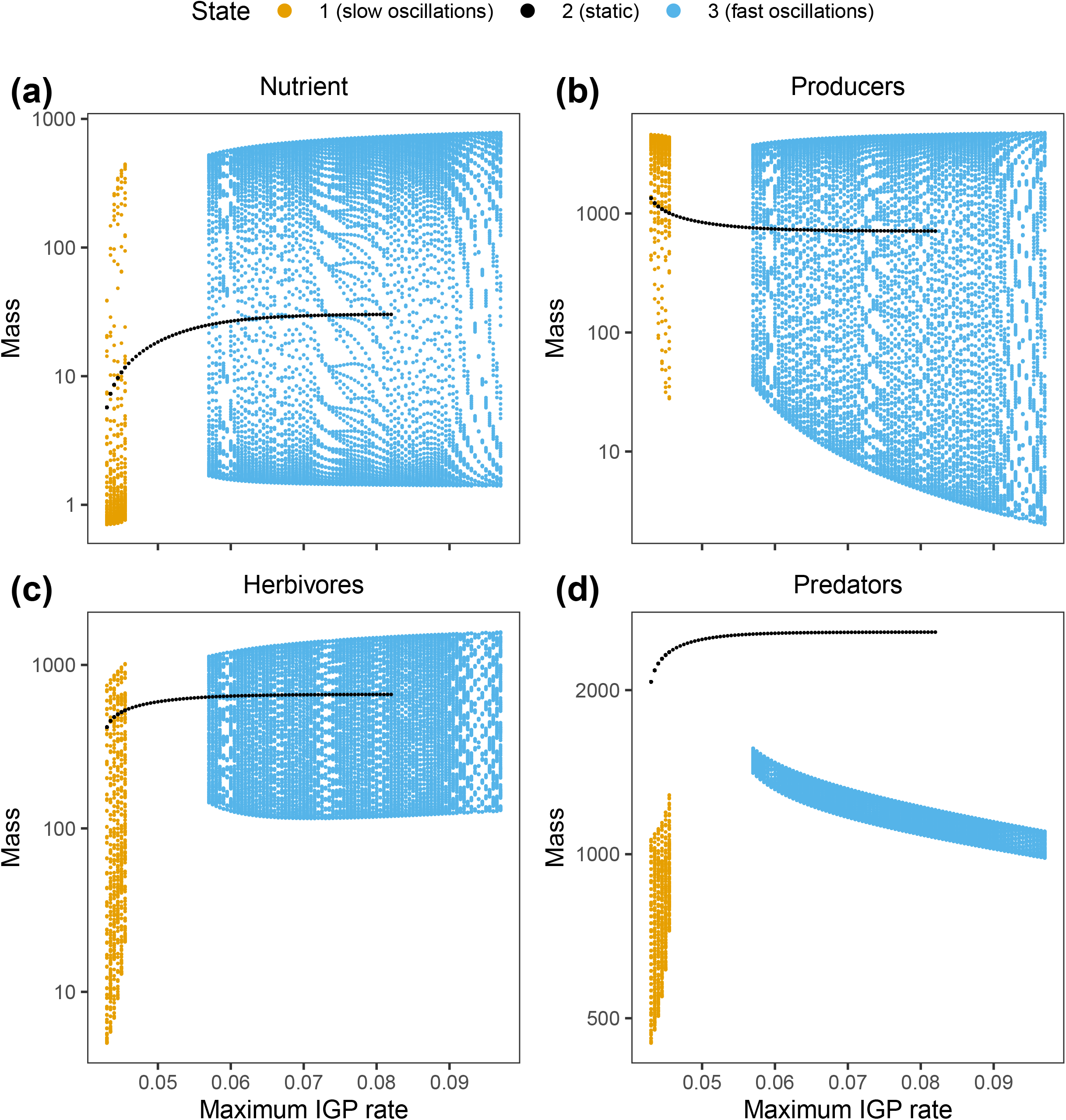
Long-run mass in µg/L of **a)** the available nutrient *N* and total biomass of **b)** the producers *A*_*i*_, **c)** the herbivores *H*_*i*_, and **d)** the predators *P*_*i*_, for simulations that differ in the maximum IGP rate 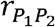 and are run with the central parameters. Many initial conditions were tested, but three initial conditions of *P*_1_ and *P*_2_ biomass were sufficient to reproduce all possible equilibrium states across 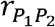 values. All outputs that result in coexistence of all six species are shown. Note this plot displays mass values at many time points. Only stable limit cycles and point equilibria were observed within the parameter ranges we studied. Also note that for most parameter combinations tested, State 1 was absent.

In State 2, the species reach a point equilibrium with no oscillations. The top predator *P*_2_ has a very high biomass, resulting in a high total predator biomass, but *P*_1_ has a very low biomass. The herbivores and producers have moderate biomasses, with *A*_2_ maintaining a somewhat high biomass but *A*_1_ maintaining a very low biomass, keeping the available nutrient at a moderately low concentration (Fig. 4 a-c, Appendix S1: Figs. S2, S3). This state occurs in a range of moderate to high maximum IGP rates: 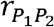 between 0.043 and 0.082 with the central parameters (Fig. 4, Appendix S1: Fig. S3), or between 0.033 and 0.116 across all parameter combinations studied (Fig. 3, Appendix S1: Fig. S1). When 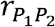 is between 0.057 and 0.082 with the central parameters, the system can also reach State 3, and it is more likely to do so the higher the initial biomass of *P*_2_ compared to *P*_1_.

Lastly, State 3 is characterized by fast, drastic oscillations across trophic levels. Trophic group and species biomasses are moderate on average, but the producers undergo downswings in which producer biomass reaches very low values while available nutrient concentration reaches high values (Fig. 2c, Fig 4a-b). This quickly cycling state occurs under higher maximum IGP rates: specifically, 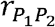 between 0.057 and 0.097 with the central parameters (Fig. 4, Appendix S1: Fig. S3) or between 0.045 and 0.267 across all parameter combinations studied (Fig. 3, Appendix S1: Fig. S1).

The effects of 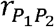 on species biomasses within states are much less pronounced than differences across states, and are detailed in Appendix S2.

### Sensitivity to nutrient enrichment and Hill exponent

The level of nutrient enrichment *n*_0_ and the Hill exponent *l* both affect the range in which the food web is found in a given qualitative state (Appendix S1: Figs. S4, S5). In general, increasing *n*_0_ somewhat constrains the range of species coexistence across 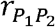 values, while decreasing *l* substantially constrains coexistence, with no coexistence possible at all when *l* =1 within the evaluated parameter sets. However, across both *n*_0_ and *l* values the main patterns of interest remain qualitatively unchanged: State 1 occurs at lower, State 2 at intermediate, and State 3 at higher 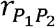 values, with State 1 being rare overall.

### Trophic interaction strengths

There is a complex relationship between the maximum IGP rate 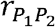 and the realized strength of IGP (eqn. 8) and other trophic interactions. The oscillating states are associated with stronger realized IGP while point equilibria are associated with weaker realized IGP (Fig. 5). In the static and quickly oscillating states (States 2 and 3), the biomasses of *A*_1_ and *P*_1_ as well as the rates at which other species consume *A*_1_ and *P*_1_ remain low compared to the biomasses and consumption of other species (Fig. 5, Appendix S1: Figs. S2, S3, S6, S7). Trophic interaction strengths are similar between States 2 and 3, with the major exception that realized IGP strength is higher in State 3 and very low in State 2 especially as 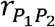 increases (Fig. 5, Appendix S1:Figs. S6 and S7). In contrast, in State 1 the distribution of trophic interaction strengths is unique. Here, most trophic interactions (rate of consumption of a nutrient or prey by a species per unit biomass of the species) are weaker than in other states, as the biomasses of most species remain low, preventing them from being consumed at a high rate (Appendix S1: Figs. S2, S3, S6, S7). On the other hand, *A*_1_ maintains a very high biomass, and the per-biomass consumption rates of *A*_1_ by the herbivores are very high. Crucially, the biomass of *P*_1_ is also much higher than in other states and, accordingly, realized IGP strength (consumption of *P*_1_ by *P*_2_) is higher than in any other state (Fig. 5, Appendix S1: Fig. S2). This is true even though the maximum IGP rate 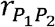 is at its lowest here, requiring that all else being equal, the consumption of *P*_1_ by *P*_2_ is lowest. Therefore, the high realized IGP strength in State 1 results from an indirect effect whereby the change in maximum IGP rate redistributes species biomasses, including an increased *P*_1_:*P*_2_ ratio (Appendix S1: Fig. S3). Consequently, realized IGP strength (the rate of consumption of *P*_1_ by *P*_2_ per unit biomass of *P*_2_) is counterintuitively high under low maximum IGP rates due to the effects of maximum IGP rate on the biomass distribution. Even so, realized IGP strength is generally highest in the cycling states and lowest in the static equilibrium state (Fig. 5).

**Figure 5:**
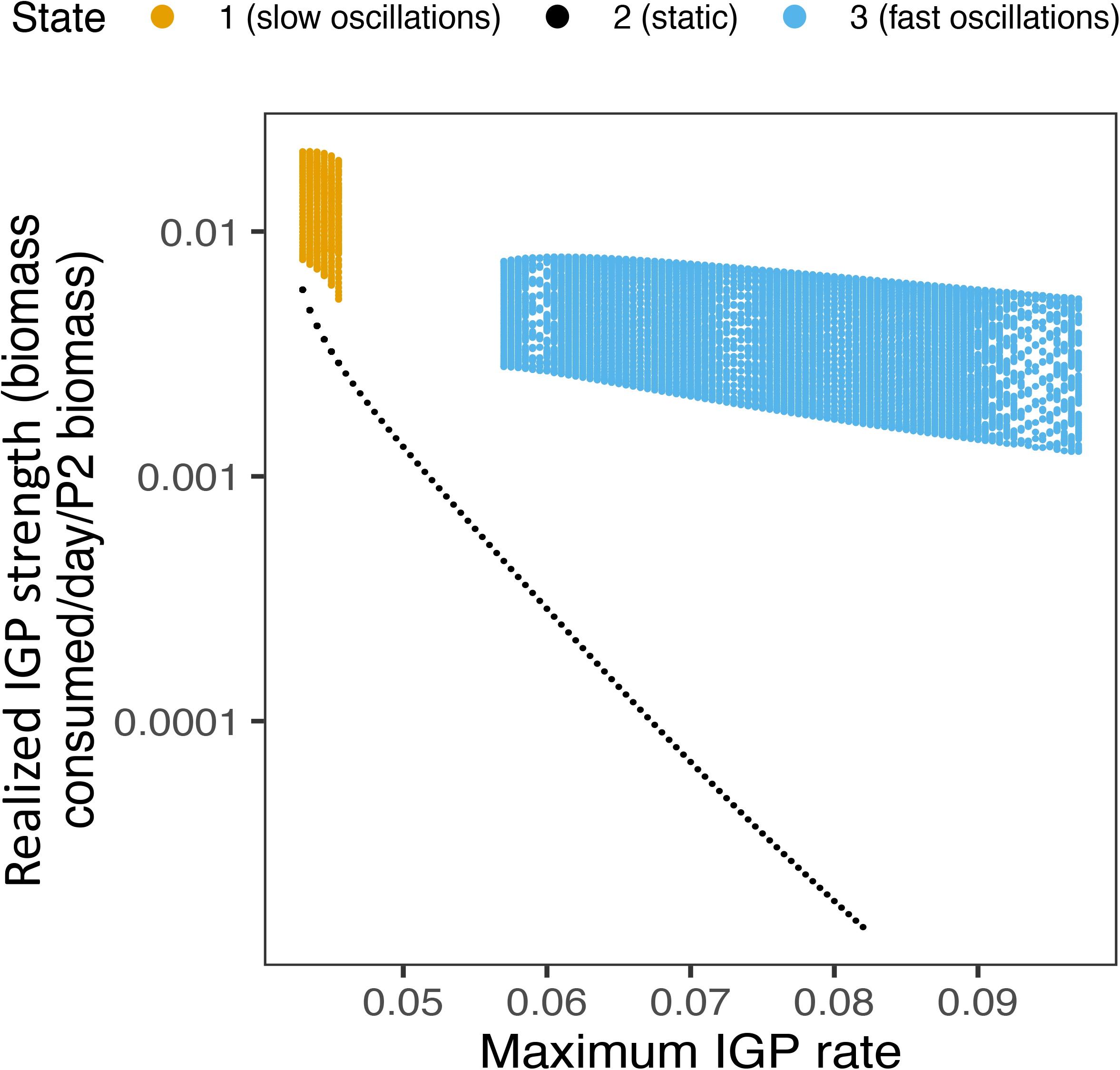
Long-run realized intraguild predation (IGP) strengths resulting from differing maximum IGP rates 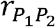. Parameters match those of Fig. 4. Realized IGP strength is the consumption rate of the smaller predator by the top predator per unit biomass of the top predator. Note this plot displays interaction strength values at many time points. Only stable limit cycles and point equilibria were observed within the parameter ranges we studied. Also note that for most parameter combinations tested, State 1 was absent.

### Causes of cycling

Analyses indicate that the strongest consumptive interaction, which differs between State 1 and 3, may induce the oscillations characterizing both of these states. In State 1, herbivory of *A*_1_ by *H*_1_ is the strongest consumptive interaction (Table 2), and only when *H*_1_ is held constant does the food web switch from State 1 to 2. In contrast, in State 3 the strongest consumptive interaction is herbivory of *A*_2_ by *H*_2_, and oscillations are eliminated only when *H*_2_ is held constant.

**Table 2:** Median strength of trophic interactions in each qualitative state. Nutrient uptake strength is measured as nutrient mass taken up per day per biomass of the producer; consumptive interaction strength is measured as biomass consumed per day per biomass of the consumer. Data are shown for the central parameters (Table 1) and 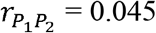 for State 1, 0.06 for State 2, and 0.08 for State 3. The strongest consumptive interaction is bolded for each state.

| **Trophic interaction** | **Median strength in State 1** | **Median strength in State 2** | **Median strength in State 3** |
| --- | --- | --- | --- |
| $N \rightarrow A_1$ | 0.0956 | 0.731 | 0.925 |
| $N \rightarrow A_2$ | 0.0438 | 0.458 | 0.652 |
| $A_1 \rightarrow H_1$ | **0.927** | 0.0231 | 0.00858 |
| $A_1 \rightarrow H_2$ | 0.627 | 0.00775 | 0.00387 |
| $A_2 \rightarrow H_1$ | 0.0124 | 0.450 | 0.132 |
| $A_2 \rightarrow H_2$ | 0.0256 | **0.475** | **0.187** |
| $H_1 \rightarrow P_1$ | 0.0112 | 0.165 | 0.130 |
| $H_1 \rightarrow P_2$ | 0.000755 | 0.0180 | 0.0129 |
| $H_2 \rightarrow P_1$ | 0.0200 | 0.0530 | 0.0450 |
| $H_2 \rightarrow P_2$ | 0.00275 | 0.0118 | 0.00919 |
| $P_1 \rightarrow P_2$ | 0.0110 | 0.000288 | 0.00386 |

## DISCUSSION

We modeled a six-species food web with four trophic levels supported by a limiting nutrient to examine cascading effects of intraguild predation (IGP) strength on the variability of species-level and trophic level biomass. The model food web converges on one of three qualitative states depending on the maximum IGP rate 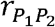 (equivalent to the inverse of handling time for the IGP interaction) and other parameters. State 1, characterized by slow oscillations, is the rarest state found only under a small range of low 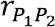 values, and is analogous to the “high-production state” in Ceulemans et al. (2019), so-called due to the relatively high primary production and producer biomass. State 2 is a point equilibrium found under low to moderate 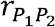. Lastly, State 3 is characterized by fast oscillations, is found under moderate to high 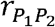, and is analogous to the “low-production state” in Ceulemans et al. (2019), so-called due to the relatively low average primary productivity. Biomass oscillations are induced by the strongest consumptive interaction, and are stronger lower in the food web such that the oscillations in producer and available nutrient mass are the most extreme. Due to food web feedback that shifts biomass distributions, realized IGP strength is counterintuitively highest under very low 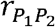 in the slowly oscillating State 1. However, sensitivity analyses reveal that State 1 is rare across parameter combinations (Fig. 3, Appendix S1: Fig. S1). Furthermore, in general both higher 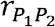 and stronger realized IGP (eqn. 8) are associated with oscillations while lower 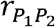 and weaker realized IGP are associated with point equilibria (Figs. 3, 5). Therefore, while the relationship between the maximum IGP rate, realized IGP strength, and biomass variability is complex in our model, the results overall support a correlation between stronger IGP in a community and biomass variability for all species.

While the generally positive observed association between IGP strength (i.e., both potential and realized IGP strength) and producer biomass variability fits with our predictions, the complexity of the association appears to echo the mixed findings of existing theory. Most of the relevant theoretical work suggests IGP is either de-stabilizing or can have various effects on stability depending on assumptions and conditions. It should be noted that previous studies, both theoretical and empirical, typically did not distinguish between potential and realized IGP strength, presumably assuming that the two were aligned. In our study, the effects of the maximum IGP rate parameter in our model are complex. Still, the complex relationship between maximum IGP rate and outcomes including biomass and realized IGP strength make sense when accounting for how the food web structure shifts. For example, a very low maximum IGP rate keeps the top predator suppressed to a low biomass, presumably due to food limitation, while allowing the small predator species to reach a very high biomass, presumably due to predation release. So even though the very low maximum IGP rate parameter requires the top predators to consume the small predators at a relatively low rate at a given density of small predators, there are so many small predators that the individual top predators still consume a relatively large amount of small predators and relatively few herbivores, leading to strong realized IGP. The absolute rate of IGP in terms of total biomass consumed per unit time (energy transfer), however, remains moderate as there are few top predators (Appendix S1: Fig. S8). Therefore, realized IGP as we define it (small predator biomass consumed per unit time per unit biomass of top predators) largely reflects the abundance of the small predator relative to the herbivore species, while the maximum IGP rate is a property of the top predator’s behavior and the absolute IGP rate integrates relative and absolute abundances.

The relationship between realized IGP strength and the central tendency of trophic level biomasses was also more complex than expected. Previous work suggested that IGP weakens trophic cascades (again, making no distinction between potential and realized IGP; Finke and Denno 2004, 2005, Straub et al. 2008), such that stronger IGP would lead to lower producer biomass. Instead, we found the opposite for both maximum IGP rate and realized IGP (Figs. 4b, 5). This result seems to be related to the balance of the energy pathways, e.g. *R*_1_ ➔ *H*_1_➔ *P*_1_ vs. *R*_2_ ➔ *H*_2_ ➔ *P*_2_. In States 2 and 3, *P*_1_ is suppressed by *P*_2_ to low biomass. However, in State 1 when the maximum IGP rate is especially low, *P*_2_ does not significantly suppress *P*_1_, so *P*_1_ maintains higher biomass than in other states. This allows *P*_1_ to suppress *H*_1_ to a lower level than in other states, freeing *R*_1_ to reach very high biomass. This is the only state in which the two predators maintain similar biomass, allowing them to more effectively partition prey (herbivore) resources and cause a stronger trophic cascade. Interestingly, the predators’ combined biomass is lowest in this state, as would be expected when (realized) IGP is high, but contrary to general expectations for when a trophic cascade is strongest. Predictions are usually that high IGP will reduce the predators’ total control of their shared prey (Finke and Denno 2005, Straub et al. 2008). However, in this case, even though the combined biomass of the predators is lower in State 1, their much more balanced biomass appears to allow them to better exploit their shared prey and cause a stronger trophic cascade by affecting both energy pathways and limiting intra-trophic-level biomass compensation. This particular finding highlights the importance of predator species evenness in shaping food web structure in addition to IGP strength.

Theoretical studies have found negative (Pimm and Lawton 1978, Holt and Polis 1997, Tanabe and Namba 2005, Hossain et al. 2020), positive (Křivan 2000), and context-dependent (McCann and Hastings 1997, Vandermeer 2006) relationships between food web stability and omnivory or IGP. There are many differences in assumptions among these models which likely contribute to the various conclusions. For example, unlike the other studies, Křivan (2000) assumed IGP is adaptive, meaning the top predator adjusts its IGP strength to maximize its population growth rate in a form of optimal foraging, which may explain why IGP provides stability in that model. Adding a fear effect, i.e. a negative relationship between predator density and prey reproductive rate, can also recover stability otherwise compromised by IGP (Hossain et al. 2020). In our model, unlike in most other models of IGP dynamics, we include an alternate herbivore species for a total of three prey species consumed by the top predator. The few other studies to do so found that adding alternate prey species to three-species IGP food webs can facilitate coexistence of the predators but can also induce unstable dynamics over a wide parameter space, which could help explain why IGP was often associated with oscillations in our model (P. Daugherty et al. 2007, Holt and Huxel 2007). Besides an additional prey species, we also included additional trophic levels and more complex functional responses for each trophic interaction in the model. Simply put, our model is more complex than the models in the aforementioned studies and could potentially reflect some real food webs more accurately, albeit at a sacrifice of generality. Compared to most terrestrial aboveground food webs, our model food web exhibits faster dynamics due to higher growth, consumption, and mortality rates, as well as stronger allometric scaling structuring these rates based on trophic level and organism size. Perhaps in food webs with different characteristics, a different relationship between IGP strength and stability may be found. The time is ripe for more empirical studies guided by IGP theory, which could bridge the gap between the various models and real food web dynamics and provide needed context for further theoretical developments.

Empirical tests of the relationship between IGP strength and biomass variability are rare, but a few relevant experiments add to the IGP-stability debate. As mentioned above, these previous studies did not distinguish between potential and realized IGP. For example, Fagan (1997) and Holyoak and Sachdev (1998), studying terrestrial arthropod and aquatic protist-bacteria food webs, respectively, concluded that the presence of omnivory was stabilizing. In contrast, other experiments using aquatic protist-bacteria food webs found that IGP increased the variability of bacterial biomass (Lawler and Morin 1993) and predator biomass (Morin 1999).

However, all of these experiments only manipulated the presence or density of omnivorous predators, and therefore were not able to separate the effects of IGP from the effects of species identity or competition between the predators (i.e., multiple food web links were added simultaneously, rather than an IGP link only). Chang and Cardinale (2020) overcame this challenge by using mesh to manipulate the degree of IGP without affecting the strength of other species interactions in an aquatic microbial food web, although they did not study biomass stability. Their paired experiment and model found that adding weak IGP to a food web decreases the biomass of the shared prey, but stronger IGP increases it. However, in our model we found a different relationship where realized IGP strength is negatively correlated with the biomass of shared prey (herbivores) within qualitative states. Again, this discrepancy could be due to the larger number of species and nonlinearities in our model which create cascading shifts in biomass throughout the food web as the IGP rate changes. It is also worth noting that the large change in realized IGP strength between the static and oscillating states in our model scarcely affects herbivore biomass, and even the correlation between IGP and herbivore biomass within states is weak. Chang et al. (2020) similarly found that when the predators in their model significantly partitioned their prey resources, as occurs in our model, the effects of IGP strength on predator biomass became weak. Indeed, the more important effects of IGP strength are the effects on biomass variability in our model, hence our focus on variability throughout the paper.

Note, we were not able to explore all solutions of our model due to the inability to solve the model analytically. However, the parameters were determined carefully to fit real data, past relevant theoretical work, and key tradeoffs found in real food webs. It is also worth noting that as most parameters are varied away from their central values, the six species coexist within a more restricted range of 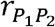. Perhaps, therefore, the central parameter values are reasonably representative of real food webs which exhibit a wide range of IGP rates. Overall, despite these caveats we are confident in the conclusion that IGP can either generate cycling behavior or maintain stability in model food webs, depending on its strength. We look forward to empirical tests of our findings and for the intuition that this model may bring to understanding the dynamics of aquatic, terrestrial, natural, and managed ecosystems.

The results of this study can be applied to biological pest control methods. When multiple predators are used as biological control agents in a single system, releasing or attracting predators which engage in low rates of IGP may have the effect of enhancing the stability of crop yields. This is particularly likely if the focal species adhere to our model assumptions, such as exhibiting relatively high birth and consumption rates and following allometric scaling, e.g. if smaller species exhibit higher birth, consumption, and mortality rates than larger species, as in belowground and planktonic systems. While our results could be applicable to some terrestrial agricultural systems, planktonic systems are also relevant to biological control. Environmental algal technologies, such as cultivating phytoplankton as a way of cleaning wastewater while producing biofuel feedstock, are promising strategies for combating multiple environmental challenges but suffer from the proliferation of zooplankton (Smith et al. 2010). Therefore biological control is promising for improving the economic feasibility of algal cultivation (Rakowski et al. 2024), and this nascent field of algal cultivation biocontrol could benefit from insight into the properties of predators that might improve the stability of algal production.

As global change further alters food webs and predation rates, the results of this study imply that increases in IGP rates could run the risk of de-stabilizing populations and key ecosystem functions such as primary production. For example, one study found that warming significantly increased IGP levels among aquatic invertebrates (Frances and McCauley 2018). However, the generality of this relationship across ecosystems remains to be determined, and there may not be reason to expect a universal de-stabilization of ecosystems mediated by changing IGP. Still, the connections explored herein shed light on the intricacies of strong indirect effects in even a relatively simple six-species food web and underline the possibility of strong cascading impacts of changing food web composition and predation rates. Fruitful avenues for future research could include empirically testing whether IGP strength is positively correlated with biomass variability across trophic levels, and improving predictions for when IGP might increase or decrease.

## ACKNOWLEDGMENTS

We would like to thank Ursula Gaedke and Matthieu Barbier for providing guidance for the model analysis. Christopher Peterson aided in writing the compact form of the model equations. We also thank members of the Farrior and Leibold groups as well as Shalene Jha for providing helpful feedback on the model analysis and earlier versions of the manuscript. The University of Texas at Austin provided financial support.

## AUTHOR CONTRIBUTIONS

CJR conceived the idea and built and analyzed the model with guidance from MAL and CEF. CJR drafted the manuscript, and all authors contributed revisions and approved the final version for publication.

