## Appendix S1 for "Strengthening intraguild predation increases temporal variability of biomass across trophic levels in model food webs"

**Appendix S1: Expanded model and simulation results for individual species and interactions.**

**Equations S1-S5:** Expanded and annotated model (see Table 1 in main text for parameter descriptions)

*Available nutrient mass*

net nutrient loading

(accounting for loss)

$\frac{dn}{dt}={\delta(n}_{0}-n)-\frac{w_{N}}{w_{C}}\sum_{i=1,2} \frac{r_{A_{i}}a_{i}n}{n+u_{i}}$	eqn. S1

nutrient uptake by *A*_1_ & *A*_2_

*Biomass of producer species:*

growth from nutrient uptake	density-independent mortality

$\frac{da_{i}}{dt}=\frac{r_{A_{i}}a_{i}n}{n+u_{i}}-\sum_{j=1,2} r_{H_{j}}h_{j}\frac{\left( \frac{a_{i}}{c_{A_{i}H_{j}}} \right)^{l}}{1+\sum_{i=1,2} \left( \frac{a_{i}}{c_{A_{i}H_{j}}} \right)^{l}}-m_{A_{i}}a_{i}$	eqn. S2

consumption by *H*_1_ & *H*_2_

*Biomass of herbivore species:*

gain from consuming *A*_1_ & *A*_2_	consumption by *P*_1_

$\frac{dh_{j}}{dt}=\sum_{i=1,2} b_{A_{i}H_{j}}r_{H_{j}}h_{j}\frac{\left( \frac{a_{i}}{c_{A_{i}H_{j}}} \right)^{l}}{1+\left( \frac{a_{1}}{c_{A_{1}H_{j}}} \right)^{l}+\left( \frac{a_{2}}{c_{A_{2}H_{j}}} \right)^{l}}-r_{P_{1}}p_{1}\frac{\left( \frac{h_{j}}{c_{H_{j}P_{1}}} \right)^{l}}{1+\left( \frac{h_{1}}{c_{H_{1}P_{1}}} \right)^{l}+\left( \frac{h_{2}}{c_{H_{2}P_{1}}} \right)^{l}}-r_{P_{2}}p_{2}\frac{\left( \frac{h_{j}}{c_{H_{j}P_{2}}} \right)^{l}}{1+\left( \frac{h_{1}}{c_{H_{1}P_{2}}} \right)^{l}+\left( \frac{h_{2}}{c_{H_{2}P_{2}}} \right)^{l}+\left( \frac{p_{1}}{c_{P_{1}P_{2}}} \right)^{l}}-m_{H_{j}}h_{j}$	eqn. S3

consumption by *P*_2_	density-independent mortality

*Biomass of P*_1_ *(smaller predator):*

gain from consuming *H*_1_ & *H*_2_	consumption by *P*_2_	density-independent mortality

$\frac{dp_{1}}{dt}=\sum_{j=1,2} b_{H_{j}P_{1}}r_{P_{1}}p_{1}\frac{\left( \frac{h_{j}}{c_{H_{j}P_{1}}} \right)^{l}}{1+\left( \frac{h_{1}}{c_{H_{1}P_{1}}} \right)^{l}+\left( \frac{h_{2}}{c_{H_{2}P_{1}}} \right)^{l}}-r_{{P_{1}P}_{2}}p_{2}\frac{\left( \frac{p_{1}}{c_{P_{1}P_{2}}} \right)^{l}}{1+\sum_{j=1,2} \left( \frac{h_{j}}{c_{H_{j}P_{2}}} \right)^{l}+\left( \frac{p_{1}}{c_{P_{1}P_{2}}} \right)^{l}}-m_{P_{1}}p_{1}$	eqn. S4

*Biomass of P*_2_ *(top predator):*

gain from consuming *H*_1_ & *H*_2_	gain from consuming *P*_1_

$\frac{dp_{2}}{dt}=\sum_{j=1,2} b_{H_{j}P_{2}}r_{P_{2}}p_{2}\frac{\left( \frac{h_{j}}{c_{H_{j}P_{2}}} \right)^{l}}{1+\left( \frac{h_{1}}{c_{H_{1}P_{2}}} \right)^{l}+\left( \frac{h_{2}}{c_{H_{2}P_{2}}} \right)^{l}+\left( \frac{p_{1}}{c_{P_{1}P_{2}}} \right)^{l}}+b_{P_{1}P_{2}}r_{{P_{1}P}_{2}}p_{2}\frac{\left( \frac{p_{1}}{c_{P_{1}P_{2}}} \right)^{l}}{1+\left( \frac{h_{1}}{c_{H_{1}P_{2}}} \right)^{l}+\left( \frac{h_{2}}{c_{H_{2}P_{2}}} \right)^{l}+\left( \frac{p_{1}}{c_{P_{1}P_{2}}} \right)^{l}}-m_{P_{2}}p_{2}$	eqn. S5

density-independent mortality

**Table S1:** Parameter constraints applied to randomized parameter sets used in the sensitivity analysis (see Methods). Constraints are based on allometric assumptions.

| **Assumption** | **Parameter constraint** |
| --- | --- |
| smaller species grow faster | $r_{A_{1}}>r_{A_{2}}$ |
| smaller species grow faster | $r_{A_{2}}>b_{A_{i}H_{j}}\cdot r_{H_{1}}$ |
| smaller species grow faster | $b_{A_{i}H_{j}}\cdot r_{H_{1}}>b_{A_{i}H_{j}}\cdot r_{H_{2}}$ |
| smaller species grow faster | $b_{A_{i}H_{j}}\cdot r_{H_{2}}>b_{H_{j}P_{k}}\cdot r_{P_{1}}$ |
| smaller species grow faster | $b_{H_{j}P_{k}}\cdot r_{P_{1}}>b_{H_{j}P_{k}}\cdot r_{P_{2}}$ |
| smaller species grow faster | $b_{H_{j}P_{k}}\cdot r_{P_{1}}>b_{P_{1}P_{2}}\cdot r_{P_{2}}$ |
| smaller species take up nutrients faster | $u_{A_{1}}<u_{A_{2}}$ |
| smaller species consume prey faster | $c_{A_{1}H_{1}}<c_{A_{1}H_{2}}$ |
| smaller species consume prey faster | $c_{A_{1}H_{1}}<c_{A_{2}H_{2}}$ |
| smaller species consume prey faster | $c_{A_{1}H_{2}}\leq c_{H_{1}P_{1}}\vee c_{A_{2}H_{2}}\leq c_{H_{1}P_{1}}$* |
| smaller species consume prey faster | $c_{H_{1}P_{1}}<c_{H_{2}P_{2}}$ |
| smaller species within trophic groups consume smaller prey faster than they consume larger prey | $c_{A_{1}H_{1}}<c_{A_{2}H_{1}}$ |
| smaller species within trophic groups consume smaller prey faster than they consume larger prey | $c_{H_{1}P_{1}}<c_{H_{2}P_{1}}$ |
| smaller species have faster baseline mortality rates | $m_{A_{1}}>m_{A_{2}}$ |
| smaller species have faster baseline mortality rates | $m_{A_{2}}>m_{H_{1}}$ |
| smaller species have faster baseline mortality rates | $m_{H_{1}}>m_{H_{2}}$ |
| smaller species have faster baseline mortality rates | $m_{H_{2}}>m_{P_{1}}$ |
| smaller species have faster baseline mortality rates | $m_{H_{2}}>m_{P_{2}}$ |

*$\vee$ indicates one or the other inequality must be true

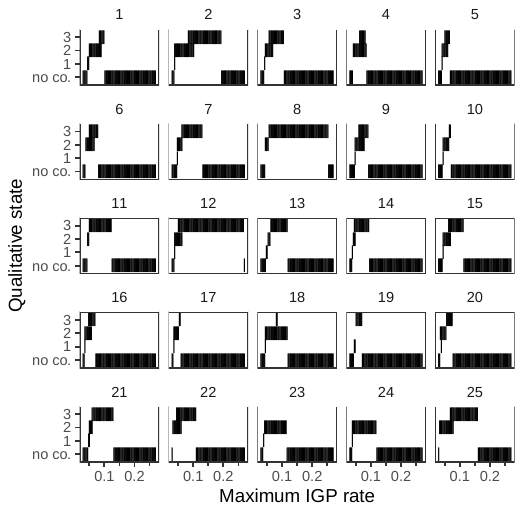

**Figure S1.** Presence of equilibrium state by $r_{P_{1}P_{2}}$ within each of 25 individual randomized parameter sets. Alternative stable states are visible as overlapping bars. “No co.” = no coexistence.

**Figure S2:** Long-run biomass of individual species – **a)** the small producer *A*_1_, **b)** the large producer *A*_2_, **c)** the small herbivore *H*_1_, **d)** the large herbivore *H*_2_, **e)** the small predator *P*_1_, and **f)** the top predator *P*_2_ – for simulations that differ in the maximum intraguild predation (IGP) rate $r_{P_{1}P_{2}}$. Simulation outputs are shown for two initial conditions of the ratio of smaller predator biomass to top predator biomass. Many other initial conditions were tested, but the system always matched one of these two alternative stable states. All simulations are run with the central parameters (Table 1) across the range of values of $r_{P_{1}P_{2}}$ that result in coexistence of all six species.

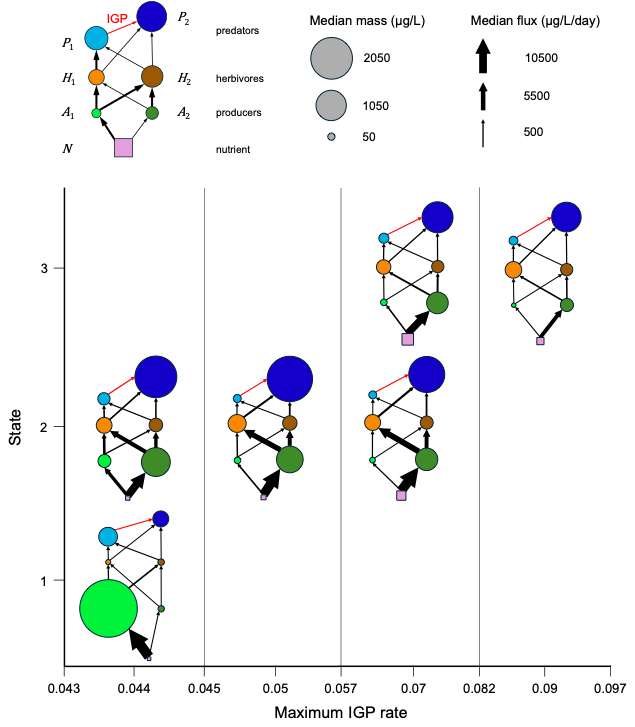

**Figure S3.** Food web diagrams representing quantitative differences in pools and fluxes across maximum IGP rate $r_{P_{1}P_{2}}$ and qualitative states. Circles represent species and have areas proportional to median C mass; squares represent the limiting nutrient and have areas proportional to median N mass on the same scale as circle area. Arrows represent mass fluxes and have widths proportional to median absolute (i.e., not relative to biomass) consumption rate of C or N for consumption of species or nutrient uptake, respectively. Results are displayed for $r_{P_{1}P_{2}}$ = {0.044, 0.05, 0.07, 0.09}. Note that the x-axis is not to scale; vertical lines and the x-axis limits represent the limits of the states displayed within those $r_{P_{1}P_{2}}$ values. All simulations are run with the central parameters (Table 1) across the range of values of $r_{P_{1}P_{2}}$ that result in coexistence of all six species.

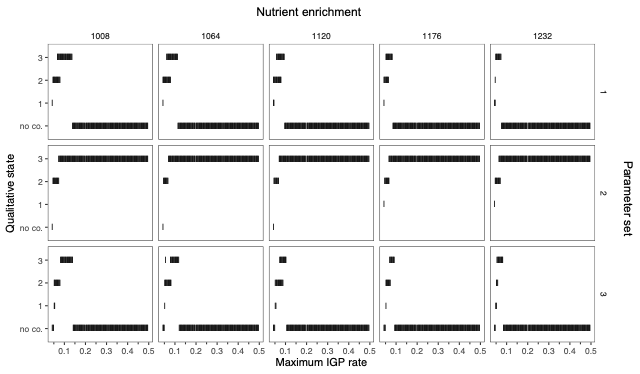

**Figure S4.** Presence of equilibrium state by $r_{P_{1}P_{2}}$ across incoming/initial nutrient concentration, i.e. nutrient enrichment, $n_{0}$, in µg*N*/L (panels from left to right), and across parameter sets (panels from top to bottom). Parameter set 1 is the central parameter set and sets 2 and 3 are randomized sets. Alternative stable states are visible as overlapping bars. “No co.” = no coexistence.

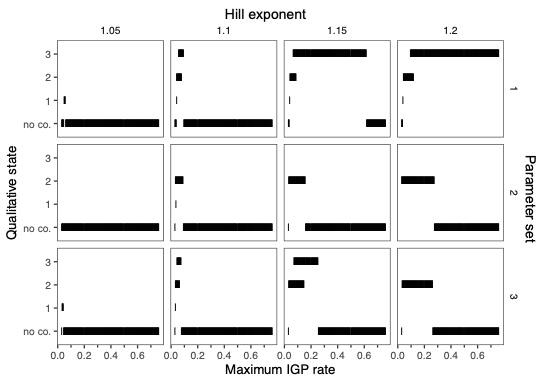

**Figure S5.** Presence of equilibrium state by $r_{P_{1}P_{2}}$ across Hill exponent, $l$, and parameter sets. Parameter set 1 is the central parameter set and sets 2 and 3 are randomized sets. Simulations were also conducted with $l$=1, but coexistence of all species was never observed under this condition for these parameter sets. Alternative stable states are visible as overlapping bars. “No co.” = no coexistence.

**Figure S6:** Long-run nutrient uptake and herbivory interaction strengths resulting from differing maximum IGP rates $r_{P_{1}P_{2}}$ (parameters match those of Fig. S1). **a)** *N* uptake by the small producer *A*_1_; **b)** *N* uptake by the large producer *A*_2_; **c)** herbivory of the small producer by the small herbivore *H*_1_; **d)** herbivory of the large producer by *H*_1_; **e)** herbivory of the small producer by the large herbivore *H*_2_; **f)** herbivory of the large producer by *H*_2_. Trophic interaction strengths are either the nutrient uptake rate per unit biomass of the producer or the consumption rate of a prey species per unit biomass of the consumer. All simulations are run with the central parameters (Table 1) across the range of values of $r_{P_{1}P_{2}}$ that result in coexistence of all six species.

**Figure S7:** Long-run non-IGP predation interaction strengths resulting from differing maximum IGP rates $r_{P_{1}P_{2}}$ (parameters match those of Figs. S1 and S2). **a)** Predation of the small herbivore *H*_1_ by the small predator *P*_1_; **b)** predation of the large herbivore *H*_2_ by *P*_1_; **c)** predation of *H*_1_ by the top predator *P*_2_; **d)** predation of *H*_2_ by *P*_2_. Trophic interaction strengths are the consumption rate of a prey species per unit biomass of the consumer. All simulations are run with the central parameters (Table 1) across the range of values of $r_{P_{1}P_{2}}$ that result in coexistence of all six species.

**Figure S8.** Long-run absolute IGP interaction strengths resulting from differing maximum IGP rates $r_{P_{1}P_{2}}$ (parameters match those of Figs. S1–S3). Absolute trophic interaction strengths are simply the consumption rate, biomass consumed/day (not relative to predator biomass), which represents the rate of energy transfer. All simulations are run with the central parameters (Table 1) across the range of values of $r_{P_{1}P_{2}}$ that result in coexistence of all six species.
