## Appendix S2 for "Strengthening intraguild predation increases temporal variability of biomass across trophic levels in model food webs"

**Appendix S2: Effects of maximum IGP rate on biomasses within the different qualitative states.**

Within the three qualitative states, there are subtler effects of changing the maximum IGP rate $r_{P_{1}P_{2}}$. In general, as $r_{P_{1}P_{2}}$ increases within a qualitative state, the available nutrient concentration and herbivore biomass increase and producer biomass decreases. Total predator biomass, on the other hand, increases with $r_{P_{1}P_{2}}$ within State 1 and 2 but decreases within State 3 (Fig. 4). However, individual species biomasses do not necessarily follow the same pattern as the total biomass of their corresponding trophic group. As $r_{P_{1}P_{2}}$ increases in State 1, *A*_2_ strongly increases even as total producer biomass decreases; and in States 1 and 2 *P*_1_ decreases while total predator biomass increases (Fig. 4, Appendix S1: Fig. S2). Oscillation amplitudes of biomass often vary less than mean biomasses as $r_{P_{1}P_{2}}$ changes. However, there is an increase in amplitude of producer biomass fluctuations as $r_{P_{1}P_{2}}$ increases (Fig. 4).
